# Early rhizosphere assembly during the onset of photosynthesis reveals inoculum-constrained succession with increased phylogenetic clustering, filtering and diversity

**DOI:** 10.64898/2026.06.25.734670

**Authors:** Rubén Chaboy-Cansado, Paula Cobeta, Gabriel Roscales, Alberto Rastrojo, Daniel Aguirre de Cárcer

## Abstract

The rhizosphere microbiome, one of the most diverse and metabolically active microbial ecosystems known, plays fundamental roles in plant health and productivity. However, the ecological dynamics occurring during the transition between germination and the establishment of the first true leaves, a developmental window associated with the onset of active photosynthesis and rapid root expansion, remain poorly understood. Here, we investigated rhizosphere microbiome assembly during the first four weeks of tomato development by sampling communities arising from seven distinct natural soil inocula twice weekly to obtain fine-scale temporal resolution.

Bacterial load, richness, evenness and phylogenetic diversity all increased significantly during plant development, indicating progressive increases in rhizosphere ecosystem complexity. In addition, diverse initial microbial communities differentially influenced both host plant development and the bacterial carrying capacity of the resulting rhizosphere ecosystem.

Although temporal effects on rhizosphere microbiome composition were significant, assembly trajectories remained strongly constrained by the initial inoculum. Temporal analysis nevertheless revealed significant taxonomic turnover despite limited global compositional restructuring. In particular, Proteobacteria and Pseudomonadaceae decreased over time, whereas Actinobacteria, Acidobacteria and Streptomycetaceae increased. However, communities did not become progressively more similar or divergent over time.

Altogether, our results indicate that early rhizosphere microbiome assembly involves rapid ecological succession within inoculum-constrained compositional trajectories, with early copiotrophic Proteobacteria progressively giving rise to more diverse and phylogenetically structured communities. These findings suggest that the first weeks of plant development may represent a critical ecological window for microbiome-based manipulation strategies in agriculture.

## INTRODUCTION

The rhizosphere, the narrow region of soil directly influenced by plant roots, harbors one of the most diverse and metabolically active microbial ecosystems known (1). Within this highly dynamic microenvironment, plants continuously shape the surrounding microbial community through the release of root exudates, mucilage, and cellular debris, thereby promoting the establishment of specific microbial populations (2, 3). In turn, rhizosphere-associated microorganisms can positively influence plant performance in multiple ways, including improving nutrient availability and phytohormone production (4), enhancing resistance to abiotic stresses (5, 6), and protecting plants against pathogens through competition or antagonistic activities (7).

The rhizosphere is characterized by intense plant–microbe and microbe–microbe interactions that strongly influence ecosystem functioning (3). Because these interactions can have profound consequences for plant health and productivity, understanding the ecological and mechanistic basis of rhizosphere community dynamics has become increasingly important. This is especially relevant in the context of the transition toward more sustainable agricultural systems, where microbiome-based approaches and products are expected to contribute substantially. Achieving this goal requires a deeper understanding of rhizosphere microbiome assembly, the interactions occurring among community members and with the host plant, and the links between microbial community composition and ecosystem function.

Li et al. (2025), working on *Astragalus mongholicus*, a medicinal leguminous species, compared germinated and non-germinated seeds and found that germination enriched bacterial taxa belonging to the genera *Pseudomonas* and *Pantoea* (8). Studies by Torres-Cortés et al. (2018) in bean and radish (9), and by Barret et al. (2015) in different Brassicaceae species (10), reported that during the transition between germination and cotyledon emergence, bacterial diversity decreased markedly and a few taxa became dominant within the community. In both studies, a strong enrichment of Gammaproteobacteria was observed, particularly members of the genera *Pseudomonas* and *Pantoea*. Moreover, the microorganisms enriched during this transition displayed typical copiotrophic traits, including rapid growth and the ability to efficiently exploit the nutrients released by seeds during germination.

The composition of the rhizosphere microbiome has been also shown to vary across the major developmental stages of plants i.e. seedling establishment, vegetative growth, and flowering. For example, Chaparro et al. (2014), studying the rhizosphere communities of *Arabidopsis thaliana*, observed clear differences in microbial community composition among developmental stages, with the vegetative stage representing a transitional state that partially overlapped both the earlier seedling stage and the later reproductive stages (11). Similarly, Moroenyane et al. (2021), working on soybean (12), also reported stage-dependent shifts in rhizosphere microbial composition and functions.

Tomato (Solanum lycopersicum) represents both one of the most economically important crop species and a widely used model for plant–microbiome research (13). Previous studies on the tomato-associated microbiome have provided important insights into microbial community composition across different plant compartments (14, 15), and the role of host genetics in shaping microbiome assembly processes (16).

Nevertheless, very limited information is available regarding the particularly important transition between cotyledon emergence and the appearance of the first true leaves, corresponding to the shift from a system largely dominated by seed-derived exudates to one increasingly dominated by photosynthesis-driven root exudation. This stage is likely to represent a particularly relevant phase for rhizosphere microbiome assembly, as it constitutes a critical transition in the establishment of the plant-associated microbial community. In addition to the potential shift in ecological filtering associated with the onset of photosynthetic activity and changes in root exudation patterns, previous studies have demonstrated that historical contingencies can strongly influence microbial community assembly trajectories since early colonizers can modify subsequent recruitment processes. Consequently, the microbial dynamics occurring during these earliest developmental stages may have long-lasting effects on the final structure and function of the rhizosphere microbiome. Understanding these processes is particularly relevant in the context of the implementation of microbiome-based strategies and synthetic microbial communities aimed at improving plant performance and sustainability in agriculture.

Here, we investigated rhizosphere microbiome assembly dynamics during the first four weeks of tomato plant development, a developmental window likely encompassing one of the periods of strongest ecological turnover during rhizosphere microbiome establishment. In tomato, this interval includes major physiological transitions such as the onset of active photosynthesis, rapid root system expansion, and the emergence of the first true leaves, with plants typically developing three to four true leaves by the end of this period. To obtain fine-scale temporal resolution of the assembly process, rhizosphere samples were collected twice per week throughout the experiment. In order to minimize the confounding effect of host genetic variation, a single tomato genotype was used. At the same time, to evaluate the reproducibility and generality of microbiome assembly patterns across different environmental microbial pools, the experiment was performed using seven distinct microbial communities originating from different soil locations.

## MATERIALS AND METHODS

### Experimental design

Seven distinct natural bacterial communities were separately inoculated onto recently germinated tomato seeds (*Solanum lycopersicum*), with five replicates each. Rhizospheres were destructively sampled every three to four days (on Mondays and Thursdays) over a four-week period. Thus, a total of 280 plants were sown, and processed along eight time-points. From these samples, bacterial load and shoot dry weight were measured, as well as rhizosphere bacterial microbiome composition.

### Natural microbial communities

The microbial fraction (MF) was extracted from various natural soils (Suppl. Table 1). Samples were collected from the upper 10 cm of soil and sieved through a clean 1 mm mesh. Each sample consisted of 120 g of soil, which was transferred into twelve sterile 50 mL conical polypropylene tubes containing 30 mL of PBS and vortexed using a Multi Reax mixer (Heidolph) at 2000 rpm for 1 h at 4 °C. To separate bacteria from soil particles, the suspensions were centrifuged for 5 min at 48 g at 5 °C in a Sigma 3-18ks centrifuge. Ten millilitres of the supernatant containing bacteria were transferred to sterile 15 mL polypropylene tubes. To remove soluble compounds from the original soil, these tubes were centrifuged at 5 °C for 30 min at 4200 g in the same centrifuge; 9 mL of the supernatant were discarded and replaced with 9 mL of cold PBS to wash the microbial pellet. The tubes were manually resuspended, centrifuged as described above, and 8.5 mL of the supernatant were removed. After this process, the remaining volumes from the twelve tubes corresponding to each sample were pooled into a single sterile 50 mL polypropylene tube. From each pool, 11.25 mL were transferred to a sterile 50 mL polypropylene tube and mixed with 3.75 mL of sterile 80 % glycerol. Finally, aliquots were prepared in sterile 0.2 mL polypropylene tubes for storage at −80 °C.

The obtained MFs were titrated before experimentation. For each MF, eight 1:10 serial dilutions were prepared in R2A medium using sterile 0.2 mL 96-well polypropylene plates. Plates were incubated at 28 °C for 48 h, after which the last wells showing growth were recorded. Bacterial titres for each MF were calculated using the Most Probable Number (MPN) approach with the MPNcalc web tool.

### Planting and inoculation

Previous studies have shown that plant genotype influences rhizosphere composition (17, 18). To avoid this confounding effect, seeds from a single genetically homogeneous tomato variety (Salinas F1, CapGen Seeds) were used. Seeds were surface-sterilized under a laminar-flow hood by immersion for 30 min in 20 mL of 50 % bleach containing 10 µL of polysorbate Tween 20 (Merck) in a sterile 50 mL polypropylene tube, rotating at 30 rpm on an LD79 Digital Test-Tube Rotator (Labinco). Seeds were then washed nine times with 20 mL of autoclaved distilled water and subsequently left for 1 h in 20 mL of autoclaved distilled water under the same agitation conditions. After sterilization, seeds were placed individually on sterile 1.5 % water-agar plates wrapped in aluminum foil and incubated at 28 °C for 48 h to germinate. Seedlings with a root of approximately 2–3 mm were selected and transplanted into sterile 40 mL polypropylene tubes containing 3 g of pre-sterilized vermiculite. Each seedling was immediately watered with 15 mL of Gamborg B5 plant culture medium (Duchefa Biochemie) pre-inoculated with the appropriate bacterial community. Each plant received an inoculum of 2 × 105 bacteria from the MFs.

Plants were maintained in sterile greenhouses under a 16 h light (24 °C) / 8 h dark (16 °C) photoperiod. Greenhouses were sterilized by washing with 70 % ethanol, sealing vents with autoclaved cotton, and exposing internal surfaces to ultraviolet light for 5 min. All remaining plants at day 18 were watered with 7 mL of sterile plant culture medium inside a laminar-flow hood.

### Plant processing

The greenhouses were opened under a laminar-flow hood to retrieve the plant-containing tubes. Shoots were separated from roots using sterile forceps, and as much vermiculite as possible adhering to the roots was removed with the same forceps. Shoots were then dried at 65 °C in an oven (JP Selecta, model 200) for 48 h to remove moisture and determine dry weight on a precision balance. Roots were placed in sterile 40 mL polypropylene tubes containing 2 mL of PBS to obtain rhizosphere microbial fractions (hereafter referred to as RMFs) and kept on ice. Once all roots were in their respective tubes, they were vortexed at 1490 rpm at 4 °C for 10 min using a Multi Reax mixer (Heidolph). After vortexing, 1 mL of each RMF was transferred to a sterile 1.5 mL polypropylene tube. Aliquots of each sample were dispensed into sterile 0.2 mL polypropylene tube strips for DNA extraction and titration.

For titration of each RMF, 1:10 serial dilutions from 1:10^2^ to 1:104 were prepared in sterile 0.2 mL polypropylene tubes prefilled with 180 µL of R2A medium. On two sterile 96-well polypropylene plates (0.2 mL wells) prefilled with 180 µL of R2A medium, six 1:10 serial dilutions were performed in duplicate, starting from 20 µL of the last dilution of each RMF prepared previously. Plates were incubated at 28 °C for 48 h, after which the last wells showing visible growth were recorded. Bacterial titres for each MF were calculated using the previously described Most Probable Number (MPN) procedure.

### Community profiling

Community DNA was extracted from each RMF using the microvolume alkaline-lysis protocol described by Bramucci et al. (2021) (19). A concentrated alkaline-lysis solution was prepared containing 21.2 mL KOH (383.2 mM), 13 mL dithiothreitol (51.86 mM) and 15.75 mL nuclease-free molecular-biology-grade water (Invitrogen). Separately, a concentrated stop solution consisting of Tris-HCl (2.54 M) was prepared. To begin the extraction, 36 µL of each sample were dispensed into the wells of a sterile 0.1 mL 96-well polypropylene plate. Twenty-seven microlitres of alkaline-lysis solution were added to each well, and within 10 min samples were vortexed briefly, centrifuged briefly, and placed at −80 °C for at least 10 min. Plates were then incubated at 55 °C for 5 min in a thermocycler, after which 27 µL of stop solution were added, followed by a quick vortex and brief centrifugation. Ninety microlitres of Mag-Bind® TotalPure NGS magnetic beads (Omega Bio-tek) were added, mixed by pipetting 10 times, and allowed to stand for 5 min. Plates were placed on a magnetic stand for 5 min and the supernatant removed, leaving DNA bound to the magnetic beads. To wash the DNA, 180 µL of 70 % ethanol were added, mixed by pipetting, left to stand for 1 min, and the supernatant removed; this washing step was repeated once. Beads were air-dried for 5 min to evaporate residual ethanol. The plate was then removed from the magnet, 30 µL of elution buffer (10 mM Tris-HCl, 1 mM EDTA) were added, mixed by pipetting several times, and left to stand for 5 min. Finally, the plate was returned to the magnet for 5 min and the supernatant containing eluted DNA was transferred to a fresh 0.1 mL 96-well plate.

The bacterial 16S rRNA gene was amplified from DNA samples using primers 799F and 1193R, which target the V5–V7 hypervariable region and including Illumina sequencing adapters separated by 0–7 random nucleotides. This strategy introduces a phase shift during sequencing and thus prevents the overlap of similar signals, substantially improving sequencing output (20). Amplifications were carried out in 25 µL reactions containing 4 µL of template DNA, 0.1 µM of each primer, 0.4 mM dNTPs, and 0.5 U of Q5 High-Fidelity DNA Polymerase (New England Biolabs). Thermocycler conditions consisted of 95 °C for 30 s, followed by 25 cycles of 95 °C for 10 s, 55 °C for 30 s, and 72 °C for 30 s, with a final extension at 72 °C for 2 min. Resulting PCR products were diluted 1:10 in 20 µL of nuclease-free water (Invitrogen) to reduce spurious amplicons and residual oligonucleotides from the first amplification. A second amplification of 10 cycles was then performed under the same conditions using 3 µL of the diluted products as template and 4 µL of 5 µM secondary primers containing Illumina i5 and i7 adapter sequences, 10-nt barcodes, and Illumina sequencing primer sites. Within each amplification batch, reactions shared a common barcode on one end and a unique barcode on the other, enabling unambiguous assignment of reads to samples.

Amplicon libraries were checked on 1.5 % agarose gels. Band intensities were visually estimated within each batch to combine approximately equimolar amounts of each sample. Equimolar libraries were pooled and concentrated with Pellet Paint® NF Co-Precipitant (Merck) according to the manufacturer’s instructions, then size-selected by horizontal agarose gel electrophoresis and purified using the Speedtools PCR Clean-Up Kit (Biotools). The final pooled amplicon libraries were sequenced on an Illumina NextSeq platform using the NextSeq1000/2000 P1 flowcell at 2 x 300.

### Sequence processing and analyses

Initial sequence processing was performed using the *DADA2* package in *R* (21), following its standard pipeline for error modelling, paired-end read merging, chimera removal, taxonomic assignment with the SILVA training set (v123) (22), and removal of residual eukaryotic sequences, including mitochondria- and chloroplast-derived reads. To ensure comparable sequencing depth, samples were rarefied to a common threshold determined by the natural break in the sequence-count distribution (23) (Suppl. Fig. 1); samples below this 24562 sequence depth threshold were excluded. For analyses based on phylogenetic metrics, amplicon sequence variants (ASVs) were mapped to the Greengenes2 99 % representative sequence set and its corresponding phylogenetic tree (24) using BLAST.

Faith’s phylogenetic diversity was calculated using the *pd* function from the *picante* package (v1.8.2) (25). Community composition was explored through nonmetric multidimensional scaling (NMDS) ordinations based on Bray-Curtis and UniFrac distance matrices using the *ordinate* function from the *phyloseq* package (v1.52.0) (26).

Linear models and analyses of variance (ANOVA) were fitted using the *lm* function in *R* and the *Anova* function from the *car* package (v3.1-3). Heteroscedasticity associated with these analyses was assessed using *bptest* from the *lmtest* package (v0.9-40), whereas robust coefficients were estimated using the *coeftest* and *vcovHC* functions from the *sandwich* package (v3.1-1). Partial correlations among the study variables were evaluated using the *pcor* function from the *ppcor* package (v1.1).

Bray-Curtis sample dissimilarities were calculated using the *vegdist* function from the *vegan* package (v2.6-10), whereas UniFrac-based dissimilarities were calculated using the *distance* function from the *phyloseq* package. These distance matrices were subsequently used to perform partial distance-based redundancy analyses (dbRDA) based on Bray-Curtis and UniFrac distances using the *capscale* function from the *vegan* package. In both cases, models were fitted by conditioning on Community type while testing for the effect of Time, and vice versa. The significance of these effects was assessed by permutation tests using the *anova* function, and adjusted R^2^ values were obtained using *RsquareAdj*, both implemented in the *vegan* package.

Associations between taxon relative abundances and Time were evaluated using the *Maaslin2* function from the *MaAsLin2* package (v1.22.0), including Community as a random effect and removing taxa below 0.01% and 10% relative abundance and prevalence thresholds, respectively, and q-values were adjusted with Benjamini-Hochberg FDR approach. To assess temporal dynamics within Community, intra-community distances were calculated from the Bray-Curtis and UniFrac distance matrices for each Time level. For each Community, the temporal slope of these distances was estimated and subsequently combined into a global statistic whose significance was evaluated through permutation testing.

The standardized effect size of the mean pairwise phylogenetic distance (MPD SES) was calculated for each sample by comparing the observed MPD against a null distribution generated using the *sample.pool* null model (999 permutations) implemented through the *ses.mpd* function of the *picante* package. To evaluate the effects of Time and Community on MPD SES, a linear mixed-effects model was employed. The significance of the model effects was assessed using Type III ANOVA.

All scripts and 16S rRNA gene dataset are available at https://github.com/microenvgen/Temporal and the European Nucleotide Archive (PRJEB114473), respectively.

## RESULTS

The rhizosphere microbiome of our experimental system was dominated by bacterial families typically associated with this type of ecosystem, such as Pseudomonadaceae, Burkholderiaceae, and Streptomycetaceae, although compositional differences were observed depending on the initially inoculated community type (hereafter, Community). Although some trends related to the effect of sampling time (hereafter, Time) could be discerned in specific communities such as B and C, overall community compositions appeared relatively stable over time (Fig. 1).

**Figure 1.**
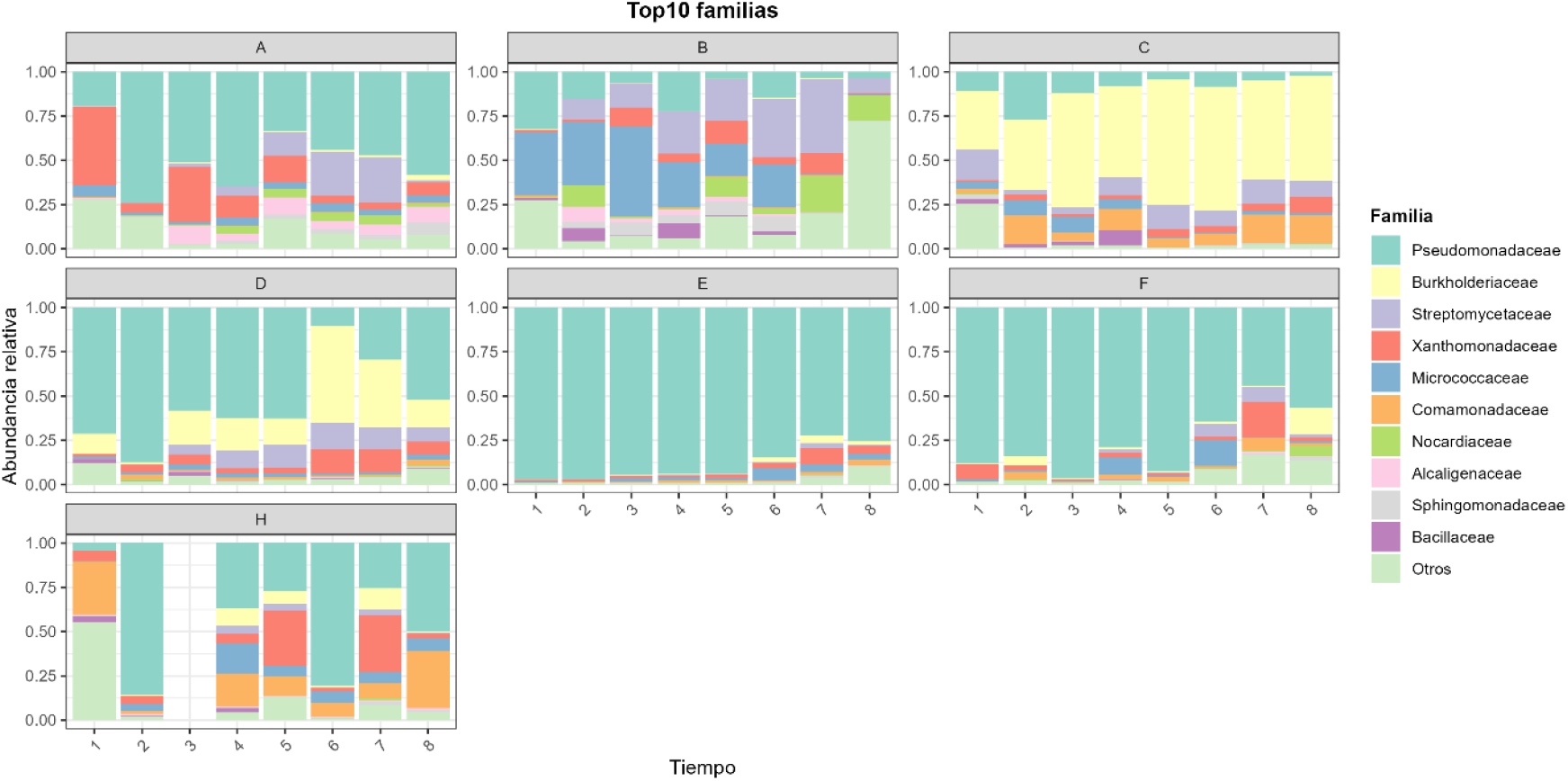
Temporal variation in the relative abundance of the 10 most abundant families across the dataset. Each panel corresponds to a different Community type and depicts the distribution of relative abundance values (Y-axis) across the different Time levels (X-axis). Owing to the filtering step applied during sample selection based on plant status, all samples from Community H were lost at the third Time level.

The representation of Load and Weight values revealed differences among Community types, and both variables increased with Time (Fig. 2A-B). Significantly, the Load-to-Weight ratio decreased over Time (Fig-2C), indicating that bacterial load increased proportionally less than plant biomass as development progressed. The effects of Time and Community on both variables were further explored using generalized linear models analyzed through Type II ANOVA. Time had a highly significant effect on Load (F_1,200_ = 82.29; P < 0.001), with a significant positive coefficient (β = 0.69; SE =SE = 0.07; t = 9.07; P < 0.001). Community also presented a significant overall effect (F_6,200_ = 2.94; P < 0.05). At the individual level, Community C showed a significant positive effect (β = 0.53; SE =SE = 0.17; t = 3.08; P < 0.001), whereas Community B presented a marginal negative effect (β = −0.39; SE =SE = 0. 2; t = −1.9; P =0.058), indicating that Community C harbored a higher bacterial load than the average, whereas Community B presented a lower load. The effect of Time further differed depending on Community type (F_6,200_ = 5.6; P < 0.001). However, at the individual level, only the interaction with Community A was significant, presenting a negative effect (β = −0.40; SE =SE = 0.18; t = -2.92; P < 0.05), thus indicating that the temporal increase in Load in this community was lower than the average.

**Figure 2.**
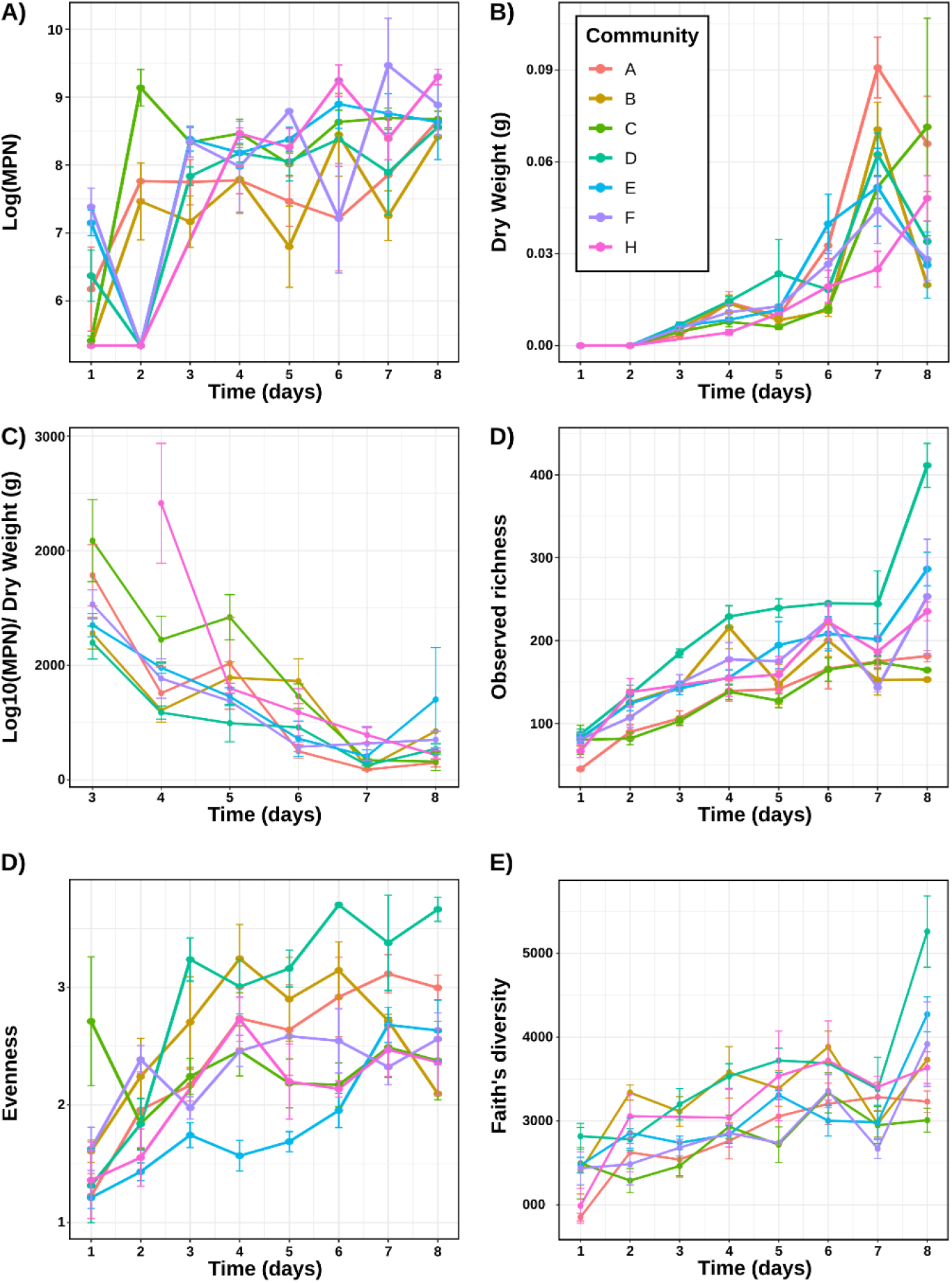
Distribution of Load (represented as the logarithmic transformation of MPN) [**A**], Weight [**B**], Load/Weight [**C**], Richness [**D**], Evenness [**E**], and Faith’s diversity [**F**] values according to Community and Time. In all cases, points represent the mean value for each Community type and bars indicate the corresponding standard deviation, with each Community represented by a different color.

The same approach was used to examine the effects of Time and Community on Weight. Again, Time presented a strong significant effect (F_1,200_ = 151.19; P < 0.001), with a significant positive coefficient (β = 0.01; SE =SE = 1.42× 10^−3^; t = 12.29; P < 0.001). By contrast, the overall effect of Community was only marginally significant (F_6,200_ = 1.87; P = 0.08), as was the interaction between Time and Community (F_6,200_ = 1.96; P = 0.07). Notably, Community A presented a significant positive effect on Weight (β = 7.48 × 10^−3^; SE =SE = 3.16 × 10^−3^; t = 2.36; P < 0.05). Furthermore, the increase in Weight over Time was more pronounced in this community, as indicated by the significant positive interaction term (β = 0.01; SE = 4.13 × 10^−3^; t = 2.93; P < 0.001), while for community F the effect was marginal and negative (β =-4.19x10^-3^ SE = 2.43 × 10^−3^; t = -1.72; P = 0,08) indicating a less pronounced increase.

The distribution of Richness and Evenness values suggested the existence of differences among Community types and, additionally, both variables appeared to increase with Time (Fig. 2). To further explore these patterns, generalized linear models analyzed through Type II ANOVA were applied to examine the effects of Time and Community on Richness and Evenness. Time exerted a highly significant effect on Richness (F_1,200_ = 212.42; P < 0.001), with a significant positive coefficient (β = 46.19; SE = 3.16; t = 14.57; P < 0.001). Community also showed a significant overall effect (F_6,200_ = 17.72; P < 0.001). In particular, Community C presented the strongest significant negative effect on Richness (β = −33.28; SE = 4.30; t = −7.72; P < 0.001), followed by Community A (β = −29.79; SE = 5.33; t = −5.58; P < 0.001), whereas Community D showed a significant positive effect (β = 61.01; SE = 9.47; t = 6.43; P < 0.001). The interaction between Time and Community also significantly affected Richness (F_6,200_ = 3.35; P < 0.05). In this regard, Community B presented a significant negative effect on Richness as Time increased (β = −19.96; SE = 4.74; t = −2.44; P < 0.05), whereas Community D exhibited a greater increase in Richness over Time compared with the remaining communities (β = 35.94; SE = 11.01; t = 3.26; P < 0.001).

Applying the same analytical approach to Evenness revealed a significant effect of Time (F_1,200_ = 77.57; P < 0.001), with a significant positive coefficient (β = 0.36; SE = 0.04; t = 8.80; P < 0.001). Community also exerted a significant overall effect on Evenness (_F6,200_ = 15.45; P < 0.001). In this case, Community E presented a significant negative effect on Evenness (β = −0.51; SE = 0.06; t = −8.20; P < 0.001), whereas Community D showed a significant positive effect (β = 0.50; SE = 0.10; t = 4.66; P < 0.001) in the same way as Community B (β = 0.32; SE = 0.11; t = 2.78; P < 0,05). The interaction between Time and Community also significantly affected Evenness (F_6,200_ = 3.37; P < 0.05). In particular, Community C presented a reduced increase in Evenness with increasing Time (β = −0.32; SE = 0.11; t = -2.76; P < 005), as did Community C albeit marginally (β = −0.13; SE = 0.07; t = -1.76; P =0.07), whereas Community D showed a comparatively faster increase in Evenness values over time (β = 0.31; SE = 0.11; t = 2,86; P < 0.001)

The phylogenetic diversity of the rhizosphere microbial communities (hereafter, Faith’s diversity) was also explored (Fig. 2). Generalized linear models analyzed through Type II ANOVA were again applied to examine the effects of Time and Community on Faith’s diversity. Both variables exerted a significant effect on this metric (F_1,200_ = 77.13; P < 0.001 and F_6,200_ = 7.40; P < 0.001, respectively). The individual coefficient associated with Time was significant and positive (β = 372.84; SE = 42.45; t = 8.78; P < 0.001). At the individual Community level, Communities C (β = −318.96; SE = 77.09; t = −4.13; P < 0.001), A (β = −234.30; SE = 79.35; t = −2.95; P < 0.05), and F (β = −172.08; SE = 87.77; t = −2.95; P = 0.05) presented marginal negative effects on Faith’s diversity, whereas Communities D (β = 480.40; SE = 117.59; t = 4.08; P < 0.001) and B (β = 264.75; SE = 93.78; t = 2.82; P < 0.05) showed significant positive effects. Although the interaction between Time and Community was not significant overall (P = 0.6),SE =

Subsequently, the effects of Richness and Diversity on Load and Weight were analyzed. Since Richness and Diversity were highly correlated (r = 0.8; P < 0.001), both the Richness variable and the fraction of Diversity independent of Richness were used in the analysis. In both cases, simple linear models were applied, and the significance of the effects was evaluated using Type II ANOVA. In the model fitted for Weight, Richness exerted a significant overall effect (F_1, 211_ = 48.31; P < 0.001), with a significant and positive individual coefficient (β = 1.52 × 10^-4^; SE = 2.19 × 10^-5^; t = 6.95; P < 0.001), indicating that an increase in Richness was associated with higher Weight. On the other hand, the fraction of Diversity independent of Richness also exerted a significant overall effect (F_1, 211_ = 6.15; P < 0.05), with a significant and negative individual coefficient (β = −9.42 × 10^-6^; SE = 3.79 × 10^-6^; t = −2.48; P < 0.05), indicating that an increase in this fraction of Diversity was associated with lower Weight. In the model fitted for Load, Richness exerted a significant overall effect (F_1, 211_ = 48.37; P < 0.001), with a significant and positive individual coefficient (β = 8.42 × 10^-4^; SE = 1.21 × 10^-4^; t = 6.95; P < 0.001), indicating that an increase in Richness was associated with higher Load values. On the other hand, the fraction of Diversity independent of Richness exerted a significant overall effect (F_1, 211_ = 11.96; P < 0.001), with a significant and negative individual coefficient (β = −7.26 × 10^-4^; SE = 2.09 × 10^-4^; t = −3.45; P < 0.001), indicating that an increase in this fraction of Diversity was associated with lower Load values.

Figure 3 (Panels A-B) depicts the NMDS ordination based on Bray-Curtis distances among the microbial communities of the studied samples. The results indicated that Community exerted a stronger effect on sample composition than Time. To further quantify this pattern, a partial distance-based redundancy analysis based on Bray-Curtis distances was performed, in which sample composition was modeled as a function of Community while controlling for the effect of Time. According to the results, Community explained 36.8% (P = 0.001) of the variation in Bray-Curtis community composition. Similarly, an NMDS ordination based on UniFrac distances, a metric that evaluates differences in microbial community composition while incorporating phylogenetic relationships, was also performed (Fig. 3, Panels C-D). Again, Community type appeared to exert the strongest effect on the separation of microbial communities, a result substantiated by a partial distance-based redundancy analysis as before indicating that Community explained 45.8% (P = 0.001) of the variation in UniFrac community composition.

**Figure 3.**
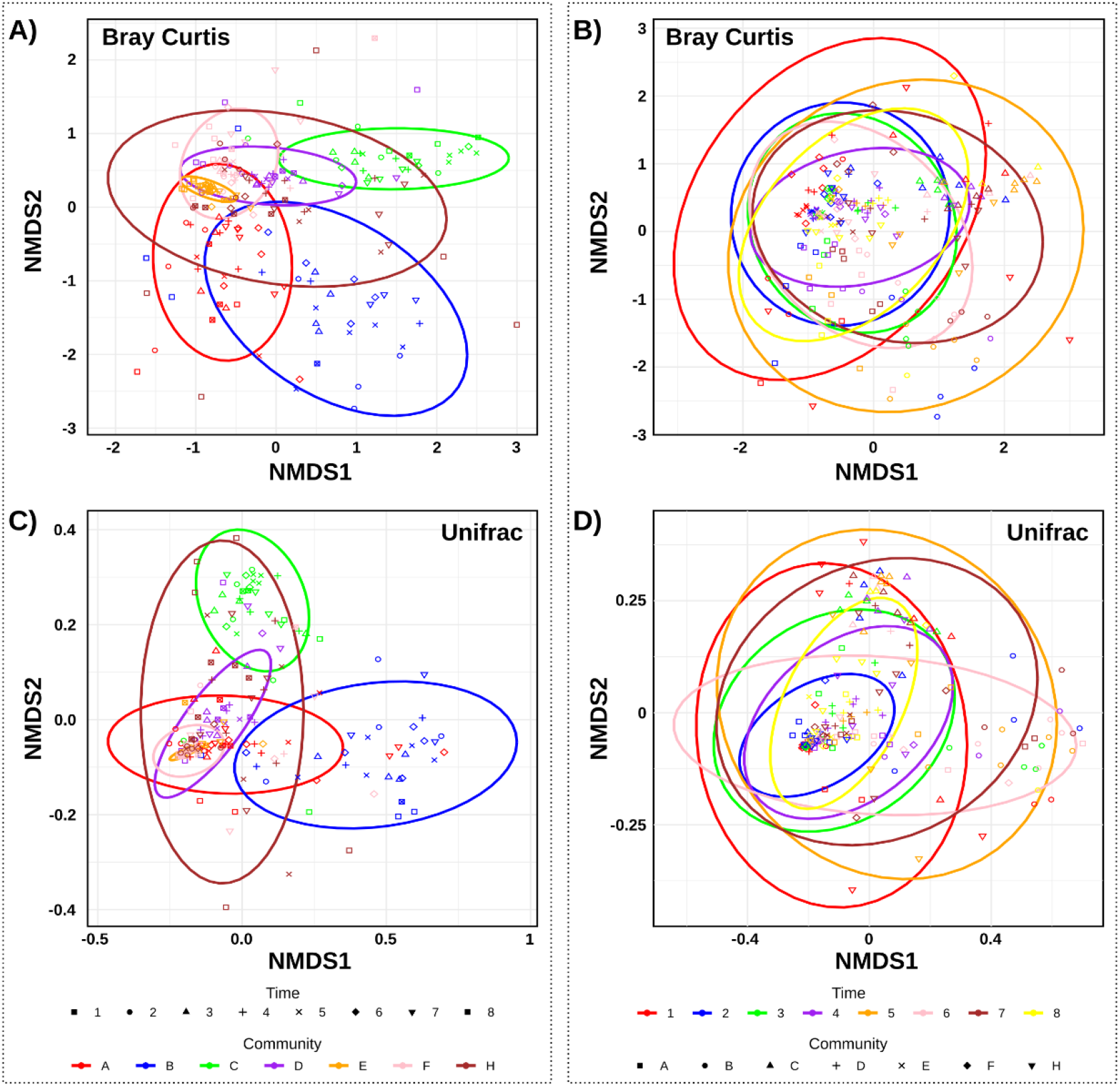
NMDS ordination based on Bray-Curtis distances (A-B) or weighted Unifrac distances (C-D) among microbial communities. **A-C;** Time is represented by shape and Community by color, with samples grouped into ellipses according to the latter variable. **B-D**, Time is represented by color and Community by shape, with samples grouped into ellipses according to Time. In both cases, ellipses represent the 95% confidence intervals for each group.

Nevertheless, when a partial distance-based redundancy analysis based on either Bray-Curtis or unifrac distances was performed, modeling sample composition as a function of Time while controlling for the effect of Community, Time was found to explain 2% and 4% (P = 0.001) of the variation in community composition, respectively. We therefore explored whether specific bacterial phyla or families exhibited significant abundance shifts over Time while controlling for the effect of Community. At the phylum level, only three phyla showed significant temporal changes in abundance (q < 0.05), with Actinobacteria and Acidobacteria increasing over Time, whereas Proteobacteria decreased in relative abundance. At the family level, five families showed a significant decrease in abundance over Time (q < 0.05), including Pseudomonadaceae, Bacillaceae, Moraxellaceae, Propionibacteriaceae, and Staphylococcaceae. By contrast, 19 families presented significant temporal increases in abundance (q < 0.05), including Streptomycetaceae, Burkholderiaceae, and Xanthomonadaceae, among others (Suppl. Table 2).

Additionally, the temporal evolution of intra-Community dissimilarity was explored using both Bray-Curtis and UniFrac distances (Suppl. Fig. 2-3). An initial inspection did not reveal a clear effect of Time on sample dissimilarity. To further assess this pattern, the slope of the change in intra-community dissimilarity across Time was estimated independently for each Community, and a global statistic was calculated as the weighted mean slope. When Bray-Curtis distances were used, the resulting statistic was −0.005 with a P value of 0.24, whereas the analysis based on UniFrac distances yielded a statistic of 0.011 with a P value of 0.93. Overall, these results did not support a significant effect of Time on intra-community dissimilarity.

Finally, to evaluate whether co-occurring taxa within individual samples were more phylogenetically related than expected by chance, the standardized effect size of the mean pairwise phylogenetic distance (MPD SES) was analyzed. MPD SES values were significantly affected by both Community (F_6, 206_ = 7.71; P < 0.001) and Time (F_1, 206_ = 137.85; P < 0.001). In particular, Time presented a significant negative coefficient (β = −1.00; SE = 0.08; t = −11.91; P < 0.001), indicating that rhizosphere communities became progressively more phylogenetically clustered over plant development.

## DISCUSSION

The rhizosphere microbiomes analyzed in this study were dominated by bacterial families typically associated with plant-associated soil ecosystems, including Pseudomonadaceae, Streptomycetaceae, Burkholderiaceae and Bacillaceae, which have been consistently reported as major components of rhizosphere communities across different plant species and soils (e.g. (11, 27-29). This overall compositional consistency suggests that, despite the use of different inoculated microbial communities, our experimental system reproduced ecologically realistic rhizosphere assemblages. At the same time, the clear compositional differences observed among inoculated communities indicate that the initial microbial pool remained a major determinant of subsequent microbiome assembly trajectories throughout the experiment.

As expected, plant weight increased throughout plant development, as did bacterial load, likely reflecting the progressive expansion of the root system and the increasing amount of carbon resources released into the rhizosphere. Nevertheless, the Load-to-Weight ratio decreased over time, indicating that bacterial load accumulated proportionally more slowly than plant biomass during plant development. Similar patterns have previously been observed in maize, where plant aging is associated with reduced exudation intensity and decreased carbon allocation to belowground compartments (30). More importantly, richness, evenness and phylogenetic diversity also increased significantly over time. These patterns cannot be explained simply as a consequence of increased bacterial load, since sequencing depth for all samples was kept constant. Instead, these results suggest that rhizosphere communities became progressively more structurally complex during the first weeks of plant development. In this context, the increase in richness may reflect the progressive expansion of ecological niche availability associated with root growth and potential changes in exudation patterns. Nonetheless, although early rhizosphere colonization has been shown to be dominated by fast-growing copiotrophs capable of rapidly exploiting the nutrient-rich conditions generated during germination and early root establishment, our results suggest that these early dominant taxa do not completely exclude the subsequent establishment of additional microbial lineages. Rather, as plant development proceeds, the rhizosphere itself may progressively become a more heterogeneous and ecologically complex environment, allowing the recruitment, persistence and coexistence of a broader range of microbial taxa and ecological strategies.

Bacterial load was positively correlated with richness, indicating that communities with higher bacterial abundance also tended to contain a larger number of detectable ASVs. At the same time, Load showed a negative correlation with phylogenetic diversity independent of richness, suggesting that the increase in bacterial abundance was not phylogenetically random but instead tended to involve the expansion of phylogenetically related lineages.

Our results indicated that the main effect on rhizosphere community composition was exerted by the identity of the initially inoculated microbial community. This result is ecologically expected, since the initial microbial pool ultimately determines which microbial lineages are available for subsequent rhizosphere assembly and which ecological interactions can potentially occur during community development. Similar strong effects of soil type or initial microbial community composition on rhizosphere microbiome assembly have been repeatedly reported when explicitly tested in different plant systems (12, 28, 31, 32). Importantly, the effect of initial community identity was not restricted to microbiome composition, as it also significantly affected bacterial Load and, albeit only marginally, plant Weight. Moreover, significant Community × Time interactions were detected for both variables. These results indicate that different microbial pools influenced both the carrying capacity of the rhizosphere ecosystem and host plant fitness.

Although temporal effects on overall community composition were statistically significant, they explained only a relatively small fraction of the total variation compared with the effect of the initial inoculated community. This result was somewhat unexpected given the major physiological transitions occurring during the studied developmental window, including the onset of active photosynthesis, rapid root growth and the emergence of the first true leaves, all of which are likely associated with substantial changes in rhizosphere resource availability. Moreover, previous studies focusing on the transition between germination and cotyledon emergence reported strong enrichments of fast-growing Proteobacteria, particularly Pseudomonas-related taxa, suggesting that early rhizosphere assembly may involve pronounced ecological turnover. However, despite these expectations, temporal trajectories in our study remained comparatively constrained, and communities did not become progressively more similar over time. These results may suggest that substantial stabilization of rhizosphere community structure occurs very early during plant development.

Nevertheless, the absence of strong global compositional restructuring did not preclude significant taxonomic turnover. In particular, Proteobacteria and Pseudomonadaceae decreased over time, whereas Actinobacteria, Acidobacteria and Streptomycetaceae increased in relative abundance, suggesting a progressive transition from early copiotroph-dominated assemblages toward more diverse and potentially more ecologically specialized rhizosphere communities.

In Arabidopsis, plant development has been shown to be associated with major shifts in root exudation profiles, with early stages enriched in sugars whereas later stages become increasingly dominated by amino acids and phenolic compounds (33). Because these latter compounds appear to exert more selective effects on microbial recruitment, the observed taxonomic shifts could potentially reflect increasing ecological filtering during rhizosphere maturation. However, since root exudates were not directly measured in our study, we cannot exclude the possibility that part of the observed succession simply reflects intrinsic microbial community maturation processes occurring independently of plant-driven exudation changes.

Interestingly, despite the observed increases in richness, evenness and phylogenetic diversity over time, rhizosphere communities became progressively more phylogenetically clustered during plant development. This pattern suggests that rhizosphere assembly was not phylogenetically random, but instead involved the progressive enrichment of specific phylogenetic lineages. One possible explanation is that rhizosphere maturation progressively increased the strength of environmental filtering exerted by the plant.

## Conclusion

Overall, our experimental system combined a high degree of environmental control with substantial ecological replication. By using a single genetically homogeneous tomato genotype grown under sterile laboratory conditions and inoculated into an initially neutral substrate, we minimized several important sources of variation typically associated with rhizosphere microbiome studies. At the same time, the inclusion of seven distinct natural microbial communities and multiple temporal replicates allowed us to evaluate the reproducibility and generality of early rhizosphere assembly dynamics across different microbial reservoirs. Importantly, our study specifically focused on the particularly narrow developmental window spanning the transition from germination to the establishment of the first true leaves, encompassing the onset of active photosynthesis and major changes in root system development. To our knowledge, although previous studies have explored either germination-associated microbiomes or broader developmental transitions, this specific early assembly window has remained comparatively underexplored at a similarly fine temporal resolution.

The identity of the initial microbial inoculum exerted a strong and expected effect on the composition of the rhizosphere communities that ultimately assembled. However, the effect of the initial microbial pool was not restricted to community composition alone, as different inoculated communities also differed in their associated bacterial loads and, to a lesser extent, in their effects on plant growth, suggesting that distinct microbial reservoirs can influence both the carrying capacity of the rhizosphere ecosystem and host plant performance. At the same time, our results indicate that the strong enrichment of fast-growing copiotrophic Proteobacteria previously reported during germination and cotyledon emergence transitions relatively rapidly toward more diverse, even and phylogenetically diverse and clustered rhizosphere communities during the following weeks of development. Although these temporal changes produced only relatively limited effects on global community composition, they were nevertheless associated with significant taxonomic turnover, suggesting the existence of rapid ecological succession processes occurring within compositionally constrained assembly trajectories. Altogether, our results suggest that the first weeks of plant development may represent a critical ecological window for microbiome-based manipulation strategies in agriculture.

## REFERENCES

1. Raaijmakers JM, Paulitz TC, Steinberg C, Alabouvette C, Moënne-Loccoz Y. The rhizosphere: a playground and battlefield for soilborne pathogens and beneficial microorganisms. Plant and Soil. 2009;321(1):341–61.

2. Bais HP, Park S-W, Weir TL, Callaway RM, Vivanco JM. How plants communicate using the underground information superhighway. Trends in Plant Science. 2004;9(1):26–32.

3. Philippot L, Raaijmakers JM, Lemanceau P, van der Putten WH. Going back to the roots: the microbial ecology of the rhizosphere. Nature Reviews Microbiology. 2013;11(11):789–99.

4. Backer R, Rokem JS, Ilangumaran G, Lamont J, Praslickova D, Ricci E, et al. Plant Growth-Promoting Rhizobacteria: Context, Mechanisms of Action, and Roadmap to Commercialization of Biostimulants for Sustainable Agriculture. Frontiers in Plant Science. 2018;Volume 9-2018.

5. Berendsen RL, Pieterse CM, Bakker PA. The rhizosphere microbiome and plant health. Trends Plant Sci. 2012;17(8):478–86.

6. Yang J, Kloepper JW, Ryu C-M. Rhizosphere bacteria help plants tolerate abiotic stress. Trends in Plant Science. 2009;14(1):1–4.

7. Mendes R, Garbeva P, Raaijmakers JM. The rhizosphere microbiome: significance of plant beneficial, plant pathogenic, and human pathogenic microorganisms. FEMS Microbiology Reviews. 2013;37(5):634–63.

8. Li D, Chen W, Luo W, Zhang H, Liu Y, Shu D, et al. Seed microbiomes promote Astragalus mongholicus seed germination through pathogen suppression and cellulose degradation. Microbiome. 2025;13(1):23.

9. Torres-Cortés G, Bonneau S, Bouchez O, Genthon C, Briand M, Jacques MA, et al. Functional Microbial Features Driving Community Assembly During Seed Germination and Emergence. Front Plant Sci. 2018;9:902.

10. Barret M, Briand M, Bonneau S, Préveaux A, Valière S, Bouchez O, et al. Emergence shapes the structure of the seed microbiota. Appl Environ Microbiol. 2015;81(4):1257–66.

11. Chaparro JM, Badri DV, Vivanco JM. Rhizosphere microbiome assemblage is affected by plant development. Isme j. 2014;8(4):790–803.

12. Moroenyane I, Tremblay J, Yergeau É. Temporal and spatial interactions modulate the soybean microbiome. FEMS Microbiol Ecol. 2021;97(1).

13. Bai Y, Kissoudis C, Yan Z, Visser RGF, van der Linden G. Plant behaviour under combined stress: tomato responses to combined salinity and pathogen stress. Plant J. 2018;93(4):781–93.

14. Dong CJ, Wang LL, Li Q, Shang QM. Bacterial communities in the rhizosphere, phyllosphere and endosphere of tomato plants. PLoS One. 2019;14(11):e0223847.

15. Ottesen AR, González Peña A, White JR, Pettengill JB, Li C, Allard S, et al. Baseline survey of the anatomical microbial ecology of an important food plant: Solanum lycopersicum (tomato). BMC Microbiol. 2013;13:114.

16. Oyserman BO, Flores SS, Griffioen T, Pan X, van der Wijk E, Pronk L, et al. Disentangling the genetic basis of rhizosphere microbiome assembly in tomato. Nat Commun. 2022;13(1):3228.

17. Rheault K, Lachance D, Morency MJ, Thiffault É, Guittonny M, Isabel N, et al. Plant Genotype Influences Physicochemical Properties of Substrate as Well as Bacterial and Fungal Assemblages in the Rhizosphere of Balsam Poplar. Front Microbiol. 2020;11:575625.

18. Yang C, Yue H, Ma Z, Feng Z, Feng H, Zhao L, et al. Influence of plant genotype and soil on the cotton rhizosphere microbiome. Front Microbiol. 2022;13:1021064.

19. Bramucci AR, Focardi A, Rinke C, Hugenholtz P, Tyson GW, Seymour JR, et al. Microvolume DNA extraction methods for microscale amplicon and metagenomic studies. ISME Communications. 2021;1(1):79.

20. Naik T, Sharda M, C P L, Virbhadra K, Pandit A. High-quality single amplicon sequencing method for illumina MiSeq platform using pool of ‘N’ (0–10) spacer-linked target specific primers without PhiX spike-in. BMC Genomics. 2023;24(1):141.

21. Callahan BJ, McMurdie PJ, Rosen MJ, Han AW, Johnson AJ, Holmes SP. DADA2: High-resolution sample inference from Illumina amplicon data. Nat Methods. 2016;13(7):581–3.

22. Quast C, Pruesse E, Yilmaz P, Gerken J, Schweer T, Yarza P, et al. The SILVA ribosomal RNA gene database project: improved data processing and web-based tools. Nucleic Acids Res. 2013;41(Database issue):D590–6.

23. Schloss PD. Rarefaction is currently the best approach to control for uneven sequencing effort in amplicon sequence analyses. mSphere. 2024;9(2):e0035423.

24. McDonald D, Jiang Y, Balaban M, Cantrell K, Zhu Q, Gonzalez A, et al. Greengenes2 unifies microbial data in a single reference tree. Nat Biotechnol. 2024;42(5):715–8.

25. Kembel SW, Cowan PD, Helmus MR, Cornwell WK, Morlon H, Ackerly DD, et al. Picante: R tools for integrating phylogenies and ecology. Bioinformatics. 2010;26(11):1463–4.

26. McMurdie PJ, Holmes S. phyloseq: an R package for reproducible interactive analysis and graphics of microbiome census data. PLoS One. 2013;8(4):e61217.

27. Lundberg DS, Lebeis SL, Paredes SH, Yourstone S, Gehring J, Malfatti S, et al. Defining the core Arabidopsis thaliana root microbiome. Nature. 2012;488(7409):86–90.

28. Edwards J, Johnson C, Santos-Medellín C, Lurie E, Podishetty NK, Bhatnagar S, et al. Structure, variation, and assembly of the root-associated microbiomes of rice. Proc Natl Acad Sci U S A. 2015;112(8):E911–20.

29. Lebreton L, Guillerm-Erckelboudt AY, Gazengel K, Linglin J, Ourry M, Glory P, et al. Temporal dynamics of bacterial and fungal communities during the infection of Brassica rapa roots by the protist Plasmodiophora brassicae. PLoS One. 2019;14(2):e0204195.

30. Pausch J, Tian J, Riederer M, Kuzyakov Y. Estimation of rhizodeposition at field scale: upscaling of a 14C labeling study. Plant and Soil. 2013;364(1):273–85.

31. Talavera-Marcos S, Gallego R, Chaboy-Cansado R, Rastrojo A, Aguirre de Cárcer D. Coupled Phylogenetic and Functional Enrichment in the Tomato Rhizosphere Microbiome. Phytobiomes Journal. 2025;9(2):151–6.

32. Schlemper TR, Leite MFA, Lucheta AR, Shimels M, Bouwmeester HJ, van Veen JA, et al. Rhizobacterial community structure differences among sorghum cultivars in different growth stages and soils. FEMS Microbiol Ecol. 2017;93(8).

33. Chaparro JM, Badri DV, Bakker MG, Sugiyama A, Manter DK, Vivanco JM. Root exudation of phytochemicals in Arabidopsis follows specific patterns that are developmentally programmed and correlate with soil microbial functions. PLoS One. 2013;8(2):e55731.

