## Supplementary for "Early rhizosphere assembly during the onset of photosynthesis reveals inoculum-constrained succession with increased phylogenetic clustering, filtering and diversity"

[1]Microbial and Environmental Genomics Group, Departamento de Biología, Universidad Autónoma de Madrid, Madrid, Spain.

| Soil | A | B | C | D | E | F | H |
|---|---|---|---|---|---|---|---|
| Origin | Guadalajara (Guadalajara) | Fuencarral (Madrid) | Valdelatas (Madrid) | Vigo (Pontevedra) | Ruiseñada (Cantabria) | Moraira (Alicante) | Algete (Madrid) |
| Soil tipe | Recreative orchard | Recreative orchard | Forest | Recreative orchard | Recreative orchard | Ruderal | Ruderal |
| Textura | sandy clay loam | sandy clay loam | silty sand | silty sand | silty sand | silty clay | sandy clay loam |
| Clay(%) | 28 | 24 | 7 | 12 | 15 | 33 | 25 |
| Sediments (%) | 14 | 6 | 10 | 21 | 24 | 26 | 10 |
| Sand (%) | 58 | 70 | 83 | 67 | 61 | 41 | 65 |
| Conductivity (µS/cm) | 330 | 1658 | 140 | 265 | 184 | 125 | 88,8 |
| pH | 7,46 | 7,21 | 5,84 | 5,46 | 6,8 | 7,74 | 7,02 |
| Organic matter (%) | 11,4 | 5,45 | 2,42 | 4,61 | 3,51 | 2,25 | 0,49 |
| Available nitrogen (mg/kg) | 5503 | 2901 | 1112 | 2606 | 2,312 | 986 | 344 |
| Available phosphorous (mg/kg) | 349 | 133 | 14,3 | 218 | 103 | < 9.80 | 13,8 |
| active lime (%) | 4,12 | 2,51 | < 0.500 | < 0.500 | < 0.500 | 7,81 | 1,13 |
| Available calcium (meq/100g) | 21,1 | 9,51 | 3,84 | 6,86 | 11,3 | 14,9 | 8,99 |
| Available magnesium (meq/100g) | 4,6 | 2,83 | 1,07 | 0,9 | 1,12 | 0,88 | 1,57 |
| Available potassium (meq/100g) | 3,46 | 4,18 | 0,41 | 0,94 | 0,61 | 0,39 | 0,26 |
| Available sodium (meq/100g) | 0,16 | 1,52 | < 0.05 | 0,1 | 0,08 | 0,09 | 0,06 |
| C/N ratio | 12 | 10,9 | 12,6 | 10,3 | 8,8 | 13,3 | 8,23 |
| Latitude | 40.64 | 40.50 | 40.54 | 42.20 | 43.36 | 38.69 | 40.61 |
| Longitude | -3.15 | -3.69 | -3.69 | -8.71 | -4.29 | 0.13 | -3.50 |
| Elevation (m) | 708 | 600 | 700 | 137 | 45 | 17 | 741 |

**Supplementary table 1. Characteristics of the soil samples used.**

**Supplementary Table 2. Bacterial families showing a significant temporal increase in abundance.**

| Family | Coeficient | q-value |
|---|---|---|
| *Streptomycetaceae* | 0,07 | < 0,001 |
| *Burkholderiaceae* | 0,05 | < 0,001 |
| *Xanthomonadaceae* | 0,02 | < 0,001 |
| *Nocardiaceae* | 0,02 | < 0,001 |
| *Sphingomonadaceae* | 0,02 | < 0,001 |
| *Rhodospirillaceae* | 0,02 | < 0,001 |
| *Rhizobiaceae* | 0,02 | < 0,001 |
| *Bradyrhizobiaceae* | 0,01 | < 0,001 |
| *Nocardioidaceae* | 0,01 | < 0,001 |
| *Hyphomicrobiaceae* | $9,64 \cdot 10^{-3}$ | < 0,001 |
| *Mycobacteriaceae* | $8,89 \cdot 10^{-3}$ | < 0,05 |
| *Caulobacteraceae* | $8,37 \cdot 10^{-3}$ | < 0,001 |
| *Flavobacteriaceae* | $8,16 \cdot 10^{-3}$ | < 0,05 |
| *Pseudonocardiaceae* | $7,98 \cdot 10^{-3}$ | < 0,001 |
| *Rhodospirillales_Incertae_Sedis* | $7,17 \cdot 10^{-3}$ | < 0,001 |
| *Chitinophagaceae* | $2,13 \cdot 10^{-3}$ | < 0,05 |
| *Xanthobacteraceae* | $1,89 \cdot 10^{-3}$ | < 0,05 |
| *SD04E11* | $1,87 \cdot 10^{-3}$ | < 0,05 |
| *Thermomonosporaceae* | $1,51 \cdot 10^{-3}$ | < 0,001 |

**SUPPLEMENTARY FIGURES.**

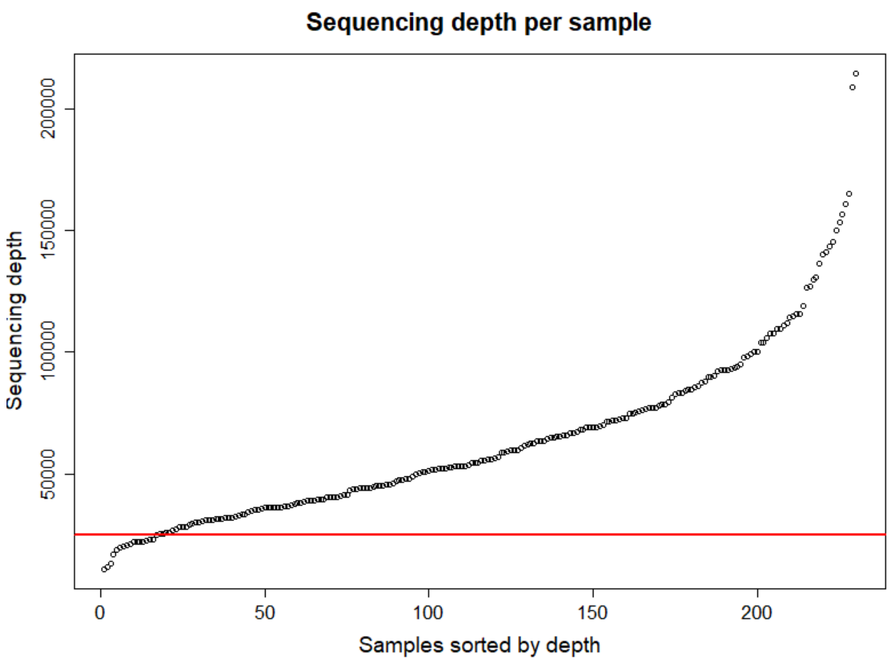


**Supplementary Figure 1. Sequencing depth per sample in the dataset.** Each point represents an individual sample (X-axis) ordered from lowest to highest sequencing depth and the corresponding number of reads per sample (Y-axis). The red line indicates the subsampling depth used (24,562 reads per sample

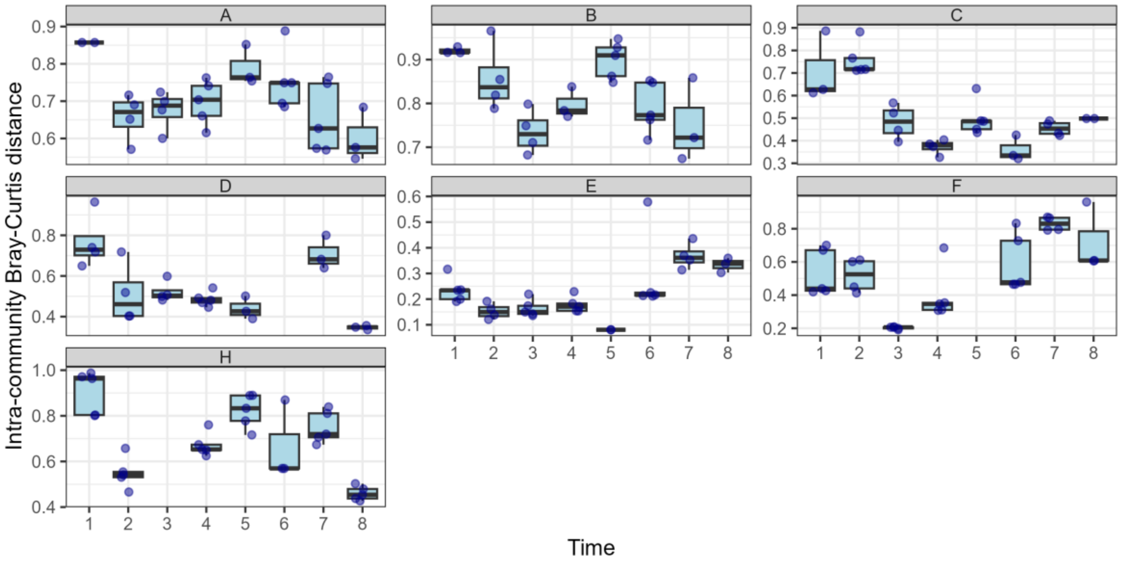


**Supplementary Figure 2.** Distribution of the temporal variation in intra-Community dissimilarity expressed as Bray-Curtis distances and represented as boxplots. Boxes represent the interquartile range, containing the central 50% of the data, whereas the line within each box indicates the median dissimilarity value among samples. Vertical whiskers extend to the maximum and minimum values excluding outliers. Each panel (A–H) corresponds to a different Community type. Again, owing to the sample filtering process, no values were available for Community H at Time 3. For those community and time combinations represented by a single replicate, it was not possible to calculate the intra-group distance, and therefore no box is shown in the plot.

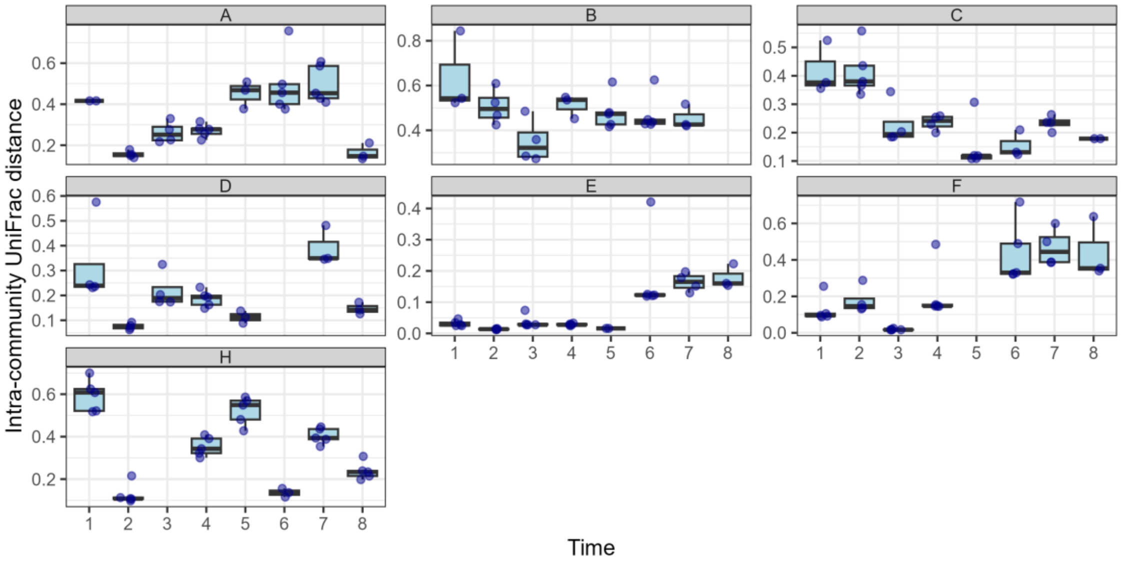


**Supplementary Figure 3.** Distribution of the temporal variation in intra-Community dissimilarity expressed as UniFrac distances and represented as boxplots. Boxes represent the interquartile range, containing the central 50% of the data, whereas the line within each box indicates the median dissimilarity value among samples. Vertical whiskers extend to the maximum and minimum values excluding outliers. Each panel (A–H) corresponds to a different Community type. Again, owing to the sample filtering process, no values were available for Community H at Time 3. For those community and time combinations represented by a single replicate, it was not possible to calculate the intra-group distance, and therefore no box is shown in the plot.
